# Engineering post-translational proofreading to discriminate non-standard amino acids

**DOI:** 10.1101/158246

**Authors:** Aditya M. Kunjapur, Devon A. Stork, Erkin Kuru, Oscar Vargas-Rodriguez, Matthieu Landon, Dieter Söll, George M. Church

## Abstract

Progress in genetic code expansion requires accurate, selective, and high-throughput detection of non-standard amino acid (NSAA) incorporation into proteins. Here, we discover how the N-end rule pathway of protein degradation applies to commonly used NSAAs. We show that several NSAAs are N-end stabilizing and demonstrate that other NSAAs can be made stabilizing by rationally engineering the N-end rule adaptor protein ClpS. We use these insights to engineer a synthetic quality control method, termed “Post-Translational Proofreading” (PTP). By implementing PTP, false positive proteins resulting from misincorporation of structurally similar standard amino acids or undesired NSAAs rapidly degrade, enabling high-accuracy discrimination of desired NSAA incorporation. We illustrate the utility of PTP during evolution of the biphenylalanine orthogonal translation system used for synthetic biocontainment. Our new OTS is more selective and confers lower escape frequencies and greater fitness in all tested biocontained strains. Our approach presents a new paradigm for molecular recognition of amino acids in target proteins.

## Introduction

Genetic code expansion broadens the structural and functional diversity of proteins and can enable synthetic biocontainment, which will be required for the 57-codon *E. coli* strain that we are constructing due to its anticipated multi-virus resistance^1^. Genetic code expansion relies on engineered orthogonal translation systems (OTSs) that use aminoacyl-tRNA synthetases (AARSs) to catalyze esterification of tRNAs to non-standard amino acids for subsequent incorporation into proteins^2^. Four primary OTS families have been developed for NSAA incorporation by suppression of amber (UAG) stop codons in targeted sequences^3,4^ (**Supplementary Fig. 1A**). However, promiscuity of engineered OTSs for standard amino acids (SAAs) and for undesired NSAAs is a major barrier to genetic code expansion. The low fidelity of several OTSs is documented, revealing that even after multiple rounds of negative selection they misacylate tRNA with SAAs that their parental variants acted upon, such as tyrosine (Tyr) and tryptophan (Trp) for engineered variants of the Tyr and Trp OTSs, respectively^5-^9^^. Furthermore, members of Tyr/Trp/Pyrrolysine (Pyl) OTS families exhibit overlap of substrate ranges, which limits their simultaneous use for potential applications involving multiple types of NSAAs^10-12^. OTS cross-talk with SAAs lowers the effectiveness of previously demonstrated synthetic biocontainment^13^ because promiscuity of the biphenylalanine (BipA) OTS^14^, which is an evolved derivative of the *Methanococcus jannaschii* Tyr OTS, can promote escape. Finally, many other NSAA applications such as protein double labelling, FRET, and antibody conjugation, require high fidelity incorporation to avoid heterogeneous protein production.

Currently, the identity of an incorporated amino acid is best determined using low-throughput protein purification and mass spectrometry. We were interested in evaluating the promiscuity of the BipA OTS and first performed higher-throughput SAA spiking experiments in minimal media lacking BipA, which suggested Tyr and Leu misincorporation (**Supplementary Fig. 1B**). We then used mass spectrometry to reveal that target peptides produced upon expression of the BipA OTS but in the absence of BipA contained 90%+ Tyr/Leu/Phe at the target site, with Q also present due to expected near-cognate suppression^15,16^ (**Supplementary Fig. 1C**). These experiments further demonstrate that SAA misincorporation is a barrier to genetic code expansion and synthetic biocontainment. To address this problem for the BipA OTS and OTSs more generally, we sought to develop a new incorporation detection system with the following design criteria: (i) the ability to controllably mask and unmask misincorporation *in vivo*; (ii) compatibility with different reporter proteins; (iii) customizability for most commonly used NSAAs. We were especially interested in associating NSAA incorporation with protein stability to accomplish these aims.

Here, we discover how the N-end rule pathway of protein degradation, a natural protein regulatory and quality control pathway conserved across prokaryotes and eukaryotes^17-19^, applies to commonly used NSAAs. We hypothesize that NSAAs may be N-end stabilizing, whereas their SAA analogs (Tyr/Phe/Trp/Leu/Lys/Arg) are known to be N-end destabilizing residues in *E. coli*, which result in protein half-lives on the timescale of minutes^18^. We test the effect of incorporation of commonly used NSAAs at the N-terminus and use our findings to develop “Post-Translational Proofreading” (PTP), which enables high-accuracy discrimination of NSAA incorporation *in vivo*. PTP is a remarkably modular, generalizable, and tunable system for specific protein recognition based on the identity of a single amino acid at the N-terminus, which is a position increasingly targeted for applications in chemical biology^20^. We subsequently show that the ability to optionally degrade proteins containing SAA misincorporation events dramatically facilitates directed evolution for selective OTSs.

## Results

### Application of the N-end rule to commonly used NSAAs

To begin investigation into how the N-end rule applies to NSAAs, we constructed a reporter consisting of a cleavable ubiquitin domain (Ub) followed by one UAG codon, a conditionally strong N-degron^21,22^, and a super-folder green fluorescent protein (sfGFP) with a C-terminal His6x-tag (Fig. 1A). We genomically integrated this reporter into a recoded *E. coli* strain devoid of UAG codons and associated release factor (C321.ΔA)^23^. The use of only one UAG codon increases assay sensitivity for promiscuity compared to the use of multi-UAG codon reporters^24^, and genomic integration of the reporter increases reproducibility by eliminating plasmid copy number effects^25^. We obtained several commonly used NSAAs and began testing with BipA (Fig. 1B). Experiments with and without BipA (BipA+ or BipA-) revealed that expression of the orthogonal 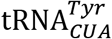 alone was responsible for a moderate amount of GFP accumulation in cells (FL/OD signal), but that expression of the BipARS together with 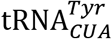 resulted in nearly equivalent signal in BipA+ or BipA-cases (Fig. 1C). Expression of an N-terminally truncated yeast Ub cleavase protein (UBP1)^26,27^ to expose the target residue at the N-terminus caused significant (∼4-fold) reduction of the BipA-signal but no significant change in the BipA+ signal. The decrease in only BipA-signal supported our hypothesis that BipA would be N-end stabilizing. BipA-signal decreased further upon overexpression of ClpS, the adaptor protein that binds peptides containing primary N-end substrates (Tyr/Phe/Trp/Leu) and delivers them to the ClpAP AAA+ protease complex for unfolding and degradation^28^, suggesting that the rate of N-end discrimination was previously limiting. ClpS overexpression may decrease substrate competition for proteins targeted for PTP because ClpS is known to inhibit other ClpA substrates such as SsrA-tagged proteins^29^.

**Fig. 1.**
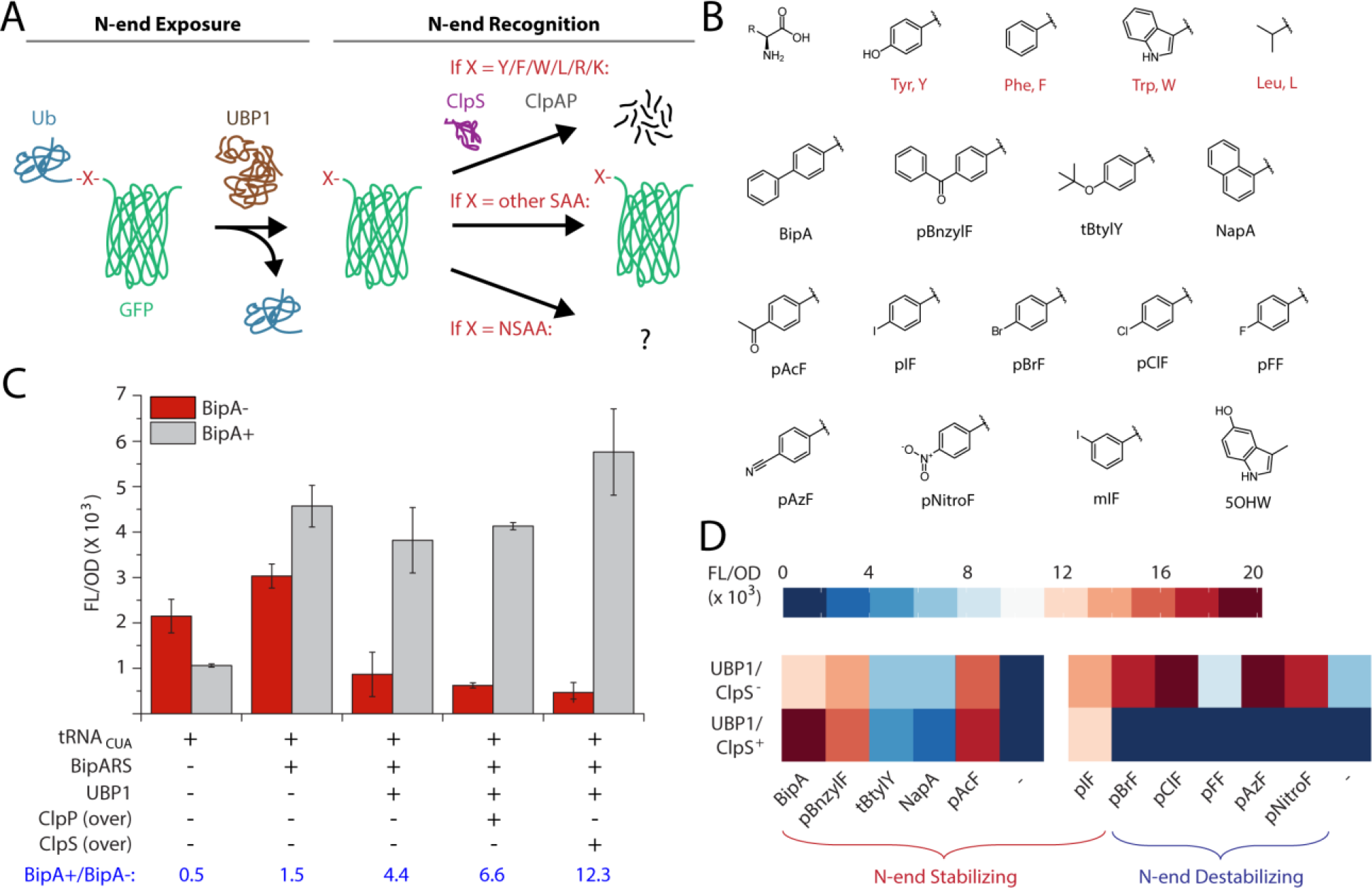
Post-Translational Proofreading (PTP) proof of concept. (A) Scheme for PTP consisting of N-end exposure and recognition steps applied to synthetic substrates. Ubiquitin (Ub) is cleaved by ubiquitin cleavase UBP1 to expose the target site as N-terminal. ClpS is the native N-recognin in *E. coli* and ClpAP forms a AAA+ protease complex for N-end rule degradation. (B) NSAAs used in this study (full chemical names in SI). (C) Incorporation assay showing fluorescence resulting from GFP expression normalized by optical density (FL/OD) in the absence/presence of BipA and expression of various OTS or N-end rule components. “Over” indicates overexpression of natively expressed components. Error bars represent SD, N=3. (D) Heatmap of FL/OD signals obtained from an NSAA panel arranged roughly in descending size from left to right without PTP in top row and with PTP in bottom row. Left panel reflects activity of BipyA OTS and right reflects pAcF OTS. Heatmap values here and elsewhere are average of N=3.

To examine how the N-end rule applies to a larger set of NSAAs, we used the bipyridinylalanine (BipyA) OTS to screen 11 NSAAs because of its low NSAA-signal (**Supplementary Fig. 2**). However, this OTS resulted in observable NSAA+ signal for only 5 out of 11 tested phenyl-NSAAs, with preference for large hydrophobic side chains at the para position of phenylalanine (Fig. 1D). Notably, NSAA+ signal for these 5 NSAAs was unaffected by PTP. The p-Acetyl-phenylalanine (pAcF) OTS was used to test incorporation of the 6 remaining NSAAs and appeared to broadly increase signal for these 6 NSAAs with PTP “Off” (ie., no expression of UBP1/ClpS). We observed marked differences in signal between PTP “Off” and “On” states based roughly on NSAA size. For p-Iodo-phenylalanine (pIF) and larger NSAAs, signal did not significantly change, and therefore pIF and larger NSAAs are N-end stabilizing. However, for p-Bromo-phenylalanine (pBrF) and other smaller or polar phenyl-NSAAs such as pAzF, signal was significantly diminished when PTP was “On” relative to when it was “Off”. The data suggest that smaller deviations from Y/F are tolerated by the ClpS binding pocket, making smaller NSAAs such as pBrF and pAzF N-end destabilizing.

### Altering N-end rule NSAA recognition using rational protein engineering

We hypothesized that we could engineer ClpS to alter N-end rule classification of these smaller NSAAs. We targeted four hydrophobic residues in the ClpS binding pocket for single point mutagenesis covering F/L/I/V (Fig. 2A). Sequence alignments of ClpS homologs across prokaryotes and eukaryotes showed conservation of these residues among related hydrophobic amino acids (**Supplementary Fig. 3**). By screening the resulting 12 single mutants in a ClpS-deficient version of our reporter strain with select NSAAs and the pAcF OTS, we identified a variant (ClpS^V65I^) that resulted in stabilization of all screened N-end phenyl NSAAs while still degrading SAAs (Fig. 2B). In addition, we identified a variant (ClpS^L32F^) that resulted in complete degradation of all but the two largest screened N-end phenyl NSAAs (**Supplementary Fig. 4**). We also attempted to distinguish tryptophanyl analogs from tryptophan using the 5OHW OTS (Fig. 2C). Although 5OHW is N-end destabilizing with ClpS, we observed that ClpS^V43I^ and ClpS^V65I^ improved discrimination of 5OHW from W (Fig. 2D). Given the desirable properties of ClpS^V65I^, we wanted to examine whether it alters N-end rule classification for SAAs. We substituted the UAG codon in our GFP reporter for codons encoding a representative panel of SAAs and found that ClpS^V65I^ affects stability of these N-end SAAs no differently than ClpS (Fig. 2E). Rational designs from our small library can precisely distinguish small modifications on a variety of chemical templates, such as NSAAs with phenyl as well as indole sidechains, showcasing the remarkably tunability of PTP. Interestingly, overexpression of either ClpS or ClpS^V65I^ lead to degradation of N-end I/V, residues that are previously shown to be only weakly N-end destabilizing *in vitro*^30^.

**Fig. 2.**
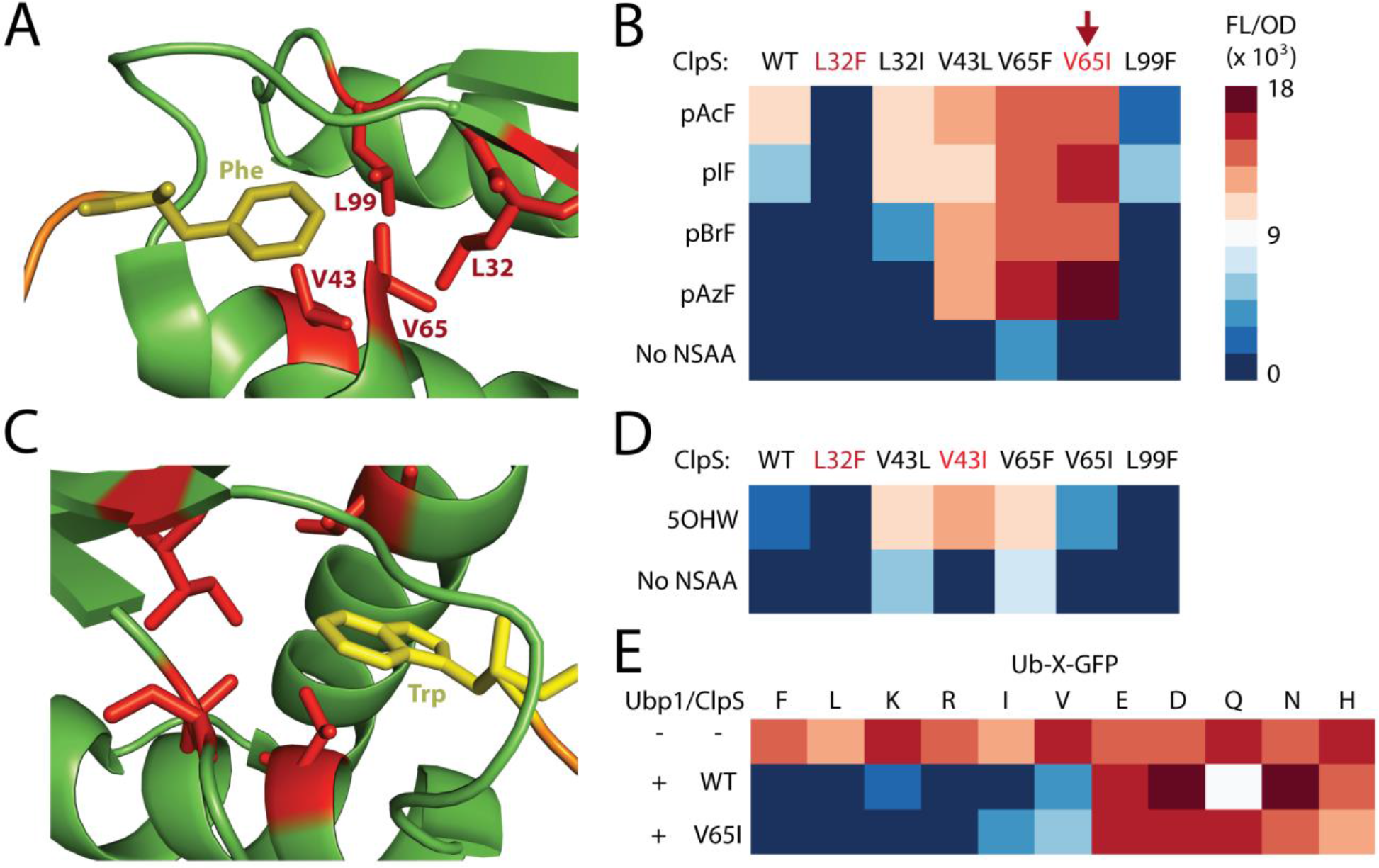
PTP tunability achieved through rational ClpS engineering. (A) Cartoon generated from crystal structure of *E. coli* ClpS binding N-end Phe peptide (PDB ID: 3O2B) showing four hydrophobic ClpS residues subjected to mutation. (B) Heatmap of FL/OD signals obtained using a ClpS-host expressing UBP1, the pAcF OTS, and variants of ClpS in the presence of different NSAAs. (C) Cartoon generated from crystal structure of *C. crescentus* ClpS binding N-end Trp peptide (PDB ID: 3GQ1). (D) FL/OD heatmap resulting from expression of UBP1, the 5OHW OTS, and ClpS variants in the presence/absence of 5OHW. Scale as in panel B. (E) FL/OD heatmap resulting from expression of UBP1/ClpS in strains with Ub-X-GFP reporter genes expressing SAAs in place of X.

### Application of PTP for selective OTS evolution

The ability of PTP to discriminate incorporation of intended NSAAs from related SAAs is useful for high-throughput screening of OTS libraries. To demonstrate this, we integrated the *UBP1-clpS^V651^* expression cassette into our ClpS-deficient reporter strain and used this strain to improve the selectivity of the parental (“WT”) BipA OTS. Previous efforts to engineer MjTyrRS variants like BipARS focused on site-directed mutagenesis on positions near the amino acid binding pocket^6,31^. To generate a novel BipARS library, we used error-prone PCR to introduce 2-4 mutations throughout the *bipARS* gene. These libraries were transformed into C321.Nend and screened with three rounds of FACS sorting: (i) positive sort for GFP+ cells in BipA+; (ii) negative sort for GFP-cells in BipA-; (iii) final positive sort for GFP+ cells in BipA+ (Fig. 3A). To decrease promiscuity against other NSAAs, we altered the negative screening stringency by varying addition of undesired NSAAs (pAcF, pAzF, tBtylY, NapA, and/or pBnzylF), which changed the profile of isolated variants (**Supplementary Fig. 5**). Retransformation of the 11 most enriched variants into C321.Ub-UAG-sfGFP (no PTP) showed that most of our variants increased BipA+ signal and decreased NSAA-signal compared to the WT OTS (Fig. 3B and **Supplementary Table 1**, Variants 1-6). Supplementation with undesired NSAAs enriched for mutants with even greater selectivity against SAAs and undesired NSAAs (Variants 4, 9-11) but also gave rise to an extremely promiscuous variant (Variant 8), suggesting that these conditions may be nearly too harsh. One mutant only isolated in higher stringencies, Variant 10, exhibited high activity on BipA and no observable activity on any other NSAAs except tBtylY, whose structure is very similar to BipA and contains the inert tert-Butyl protecting group (Fig. 3B). SDS-PAGE of Ub-X-GFP resulting from the Variant 10 OTS after expression and affinity purification showed no observable BipA-protein production in contrast to WT BipA OTS, which shows a distinct BipA-band (**Supplementary Fig. 6A**). Furthermore, mass spectrometry confirmed site-specific BipA incorporation (**Supplementary Fig. 6B-D**).

**Fig. 3.**
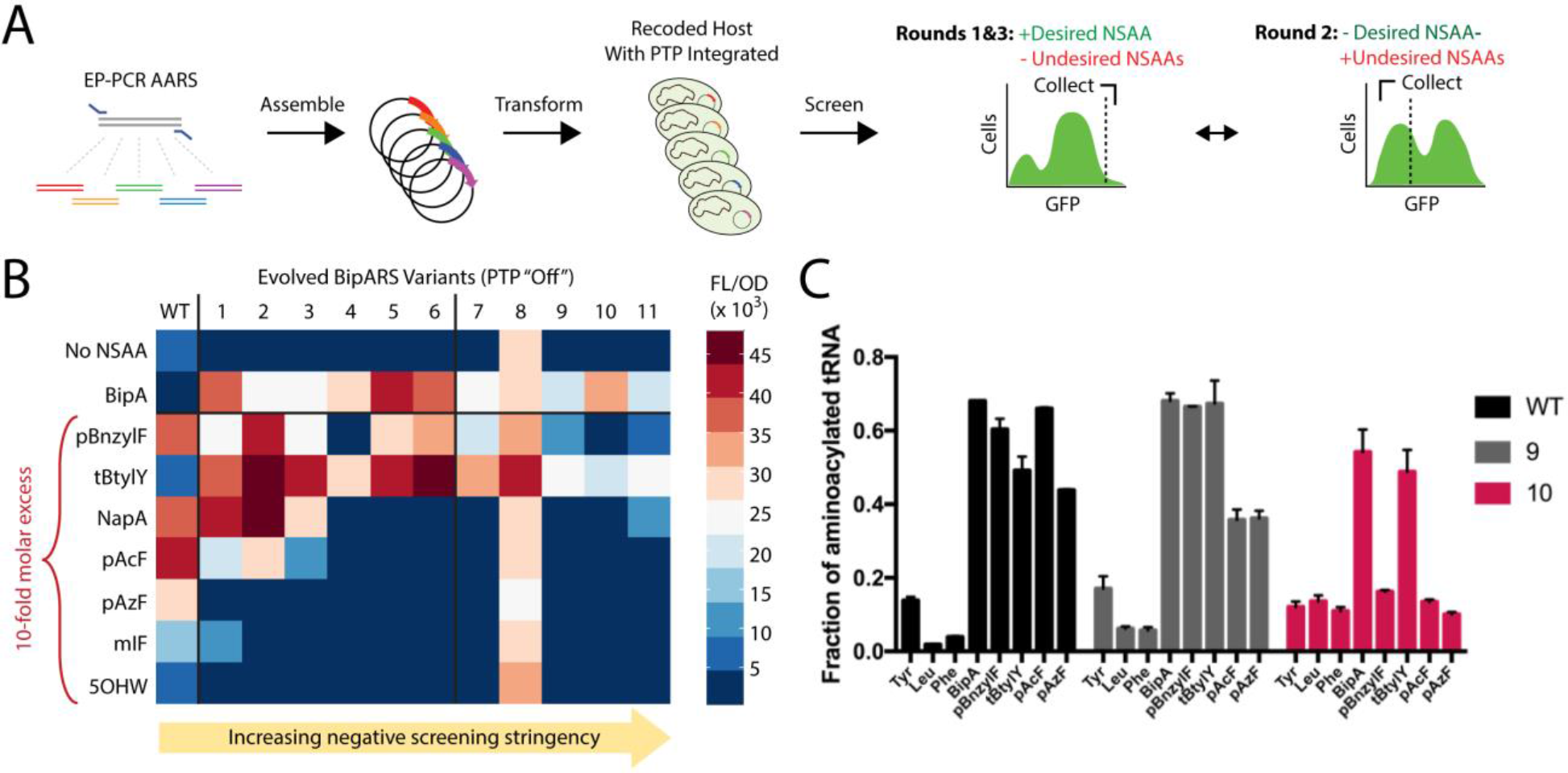
Selective BipA OTS evolution using PTP. (A) FACS evolution scheme with EP-PCR AARS libraries transformed into hosts with PTP (using ClpS^V65I^) genomically integrated before 3 sorting rounds. (B) Evaluation of most enriched evolved BipARS variants in clean backgrounds on a panel of NSAAs ([BipA] = 100 μM, [rest] = 1 mM, which are their standard concentrations). (C) *In vitro* amino acid substrate specificity profile of BipA OTS variants. Error bars = SD, N=3.

We discovered spontaneous tRNA mutations in our most selective variants, such as 4, 9, and 10, perhaps because of our use of a MutS-deficient host (**Supplementary Fig. 7A**). When we reverted these tRNA mutations, each corresponding BipA OTS became more promiscuous (**Supplementary Fig. 7B**), suggesting that observed tRNA mutations increase selectivity. The G51 position (G50 in *E. coli* nomenclature) mutated in tRNA Variant 10 is the most significant base pair in determining acylated tRNA binding affinity to elongation factor Tu (EF-Tu), which influences incorporation selectivity downstream of the AARS^32,33^. To more rigorously assess OTS selectivity, we purified AARS and tRNA for the WT, Variant 9, and Variant 10 OTSs. The observed *in vitro* substrate specificity as determined by tRNA aminoacylation is in excellent agreement with our *in vivo* assays (Fig. 3C), and the data suggests that AARS and tRNA variants each contribute to selectivity improvements (**Supplementary Fig. 7C-D**). The Variant 10 OTS exhibited the highest selectivity for BipA and was chosen for subsequent applications.

### Demonstration of enhanced biocontainment using a more selective evolved OTS

To demonstrate the utility of a more selective OTS for synthetic biocontainment, we substituted the WT BipA OTS construct previously used in three biocontained strains that exhibit observable escape frequencies with plasmids containing either the WT or Variant 10 OTS. These three biocontained strains (*adk.d6, tyrS.d8*, and *adk.d6/tyrS.d8*) harbor computational redesigns of two essential genes (*adk* and *tyrS*) to make their stability dependent on BipA^13^. We monitored escape frequencies on non-permissive media for seven days and observed lower escape frequencies for strains containing the Variant 10 OTS at all measured time-points (Fig. 4A-C, **Supplementary Fig. 8**). The difference in escape frequency was most apparent for the *adk.d6/tyrS.d8* strain, which exhibited a 7-day escape frequency of 7.4 × 10^−9^, a value more than two orders of magnitude lower than observed for any C321.ΔA-derived strain containing only two altered genes. Furthermore, the fitness of all three strains improved with the Variant 10 OTS, with doubling time decreasing by nearly 2-fold (Fig. 4D). Finally, Variant 10 also delayed onset of growth of *adk.d6/tyrS.d8* on non-cognate NSAAs (**Supplementary Table 2**). We expect these benefits to carry over to all strains which employ Variant 10 over WT OTS.

**Fig. 4.**
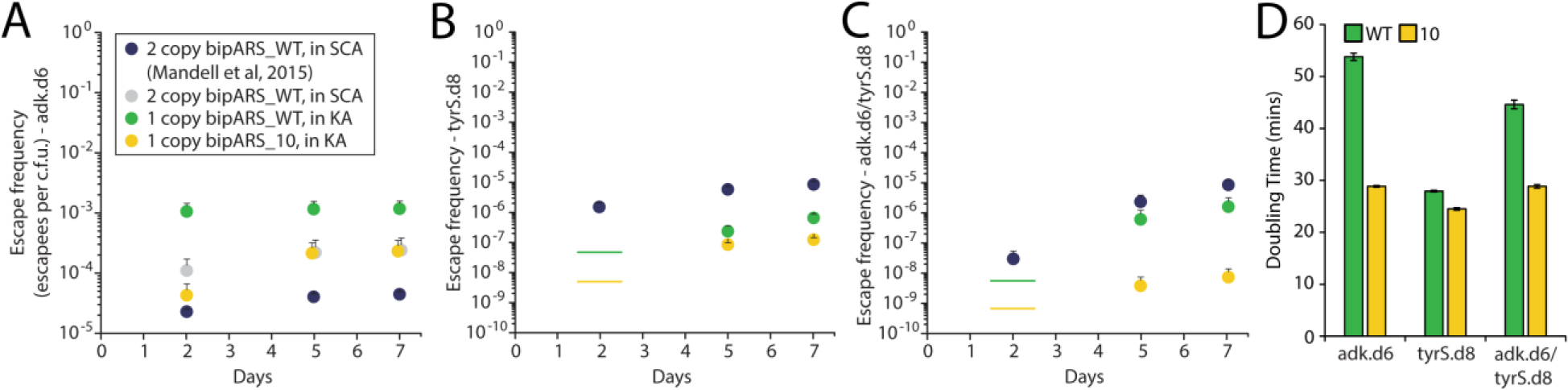
The selective evolved BipA OTS lowers escape rates and improves fitness for biocontained *E. coli* strains. (A) Escape frequencies over time for *adk.d6* strains transformed with constructs indicated in legend. Navy circles represent previously published data. Gray circles for adk.d6 represent repeat of previously published data (Ref. 13). Green and yellow circles are the most relevant constructs to compare for this study. KA: Kanamycin+Arabinose. SCA: SDS+Chloramphenicol+Arabinose. Error bars in A-C represent SEM, N=3. (B) Escape frequencies over time for *tyrS.d8* strains. Lines represent assay detection limit in cases where no colonies were observed. (C) Escape frequencies over time for *adk.d6/tyrS.d8* strains. (D) Doubling time for biocontained strains with WT or Variant 10 OTS. Error bars = SD, N=3.

## Discussion

We have demonstrated for the first time how the N-end rule applies to commonly used NSAAs, and we have subsequently engineered it to alter N-end rule classification of these molecules. We harnessed these findings to develop PTP, which significantly reduces most false positive protein expression and therefore dramatically improves the ability to determine and increase the selectivity of OTSs used for NSAA incorporation. Furthermore, the capability of PTP to distinguish among NSAAs will facilitate future efforts to simultaneously harness multiple NSAAs. We validated PTP during evolution of the BipA OTS, which resulted in greater selectivity *in vivo* and *in vitro*. Our evolved BipA OTS enhanced biocontainment efficacy and strain fitness in all tested biocontained strains.

In addition to providing a new paradigm for OTS evaluation and evolution, PTP can be transformative for applications in which the identity of a single amino acid is critical, such as screening of natural synthetases for NSAA acceptance, sense codon reassignment, post-translational modifications, and for industrial uses where purity is extremely important, such as NSAA-containing biologics. PTP may also find use in translational regulation and as an orthogonal biocontainment strategy. Given that all 20 SAAs are known to be N-end destabilizing under certain contexts^34^, conditionally expressed components could be transferred across organisms to dramatically alter the set of N-end destabilizing SAAs for a particular application.

## Online Methods

### Plasmids and plasmid construction

Two copies of orthogonal MjTyrRS-derived AARSs and 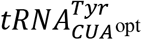 were kindly provided in pEVOL plasmids by Dr. Peter Schultz (Scripps Institute)^6^. AARSs used in this study were the following: BipARS^14^, BipyARS^14^, pAcFRS^35^, pAzFRS^36^, and NapARS^37^ The pEVOL plasmids were maintained using chloramphenicol. Original plasmids harboring two AARS copies were used for synthetase promiscuity comparison experiments (Fig. 1D, **Supplementary Fig. 2**). For generation and characterization of synthetase variants, plasmids harboring only one AARS copy under inducible expression were constructed using Gibson assembly^38^. The ScWRS-R3-13 AARS was synthesized as codon-optimized for expression in *E. coli* and cloned into the pEVOL plasmid along with its associated tRNA^39,40^. In all cases, tRNA is constitutively expressed and AARS expression is either arabinose inducible or constitutive.

An N-terminally truncated form of the UBP1 gene from *Saccharomyces cerevisiae*^26,21^ (ScUBP1^trunc^ or simply UBP1) was synthesized as codon-optimized for expression in *E. coli* and cloned into the pZE21 vector (Kanamycin resistance, ColE1 origin, TET promoter) (Expressys). The *E. coli* genes *clpS* and *clpP* were PCR amplified from *E. coli* MG1655 and cloned into artificial operons downstream of the UBP1 gene in the pZE21 vector using Gibson assembly. Artificial operons were created by inserting the following RBS sequence between the UBP1 and clp genes: TAATAAAAGGAGATATACC. This RBS was originally designed using the RBS calculator^41^ and previously validated in the context of another artificial operon^42^. Rational engineering of ClpS variants was performed by dividing the *clpS* gene into two amplicons where the second amplicon contained a degenerate NTC or NTT sequence in the oligo corresponding to each codon of interest. The four initial positions of interest in the *clpS* gene correspond to amino acids 32, 43, 65, and 99. In each case, Gibson assembly was used to ligate both amplicons and the backbone plasmid. The pZE/UBP1/ClpS and pZE/UBP1/ClpS_V65I plasmids are available from Addgene.

Three reporter constructs were initially cloned into pZE21 vectors before use as templates for PCR amplification and genomic integration. The first of these consists of a Ubiquitin-*-LFVQEL-sfGFP-His6x fusion (“Ub-UAG-sfGFP”) downstream of the TET promoter. The second has an additional UAG codon internal to the sfGFP at position Y151* (“Ub-UAG-sfGFP_151UAG”). The third has an ATG codon (encoding methionine) in place of the first UAG (“Ub-M-sfGFP_151UAG”).

### Strains and strain engineering

*E. coli* strain C321.ΔA (CP006698.1), which was previously engineered to be devoid of UAG codons and RF1, was the starting strain used for this study^23^. The TET promoter and Ub-UAG-sfGFP expression cassette was genomically integrated using λ Red recombineering^43^ and *tolC* negative selection using Colicin El^44,45^. This resulted in strain C321.Ub-UAG-sfGFP. Please see **Supplementary Table 3** for sequences of key constructs such as the reporter construct. Multiplex automatable genome engineering (MAGE)^46^ was used to inactivate the endogenous *mutS* and *clpS* genes when needed and to add or remove UAG codons in the integrated reporter. For MAGE, saturated overnight cultures were diluted 100-fold into 3 mL LB^L^ containing appropriate antibiotics and grown at 34 °C until mid-log. The integrated Lambda Red cassette in C321. ΔA derived strains was induced in a shaking water bath (42 °C, 300 rpm, 15 minutes), followed by cooling culture tubes on ice for at least two minutes. These cells were made electrocompetent at 4 °C by pelleting 1 mL of culture (16,000 rcf, 20 seconds) and washing twice with 1 mL ice cold deionized water (dH2O). Electrocompetent pellets were resuspended in 50 μL of dH2O containing the desired DNA. For MAGE oligonucleotides, 5 μM of each oligonucleotide was used. Please see **Supplementary Table 4** for a list of all oligonucleotides used in this study. For integration of dsDNA cassettes, 50 ng was used. Allele-specific colony PCR was used to identify desired colonies resulting from MAGE as previously described^47^. Colony PCR was performed using Kapa 2G Fast HotStart ReadyMix following manufacturer protocols and Sanger sequencing was performed by Genewiz to verify strain engineering. The strains C321.Ub-UAG-sfGFP, C321.Ub-UAG-sfGFP_UAG151, and C321.ΔClpS.Ub-UAG-sfGFP are available from Addgene. Ub-X-GFP reporters containing codons encoding SAAs in place of UAG were generated from Ub-UAG-GFP by PCR and Gibson assembly, and they were subsequently cloned into the pOSIP-TT vector for Clonetegration (one-step cloning and chromosomal integration) into NEB5α strains^48^. The UBP1/clpS_V65I operon was also placed under weak constitutive expression and integrated into C321.ΔClpS.Ub-UAG-sfGFP using Clonetegration. This strain (C321.Nend) was used as the host for FACS experiments.

### Culture Conditions

Cultures for general culturing, for experiments in Figure 1, for FACS screening, and for biocontainment escape assays were grown in LB-Lennox medium (LB^L^: 10 g/L bacto tryptone, 5 g/L sodium chloride, 5 g/L yeast extract). Cultures for all other experiments in Figures 2 and 3 were grown in 2X YT medium (2XYT: 16 g/L bacto tryptone, 10 g/L bacto yeast extract, 5 g/L sodium chloride) given improved observed final culture densities compared to LB^L^ upon expression of ClpS variants. Unless otherwise indicated, all cultures were grown in biological triplicate in 96-well deep-well plates in 300 μL culture volumes at 34 °C and 400 rpm.

### Chemicals

NSAAs and SAAs used in this study were purchased from PepTech Corporation, Sigma Aldrich, Santa Cruz Biotechnology, Bachem, and Toronto Research Chemicals. The following amino acids were purchased: L-4,4-Biphenylalanine (BipA), L-4-Benzoylphenylalanine (pBnzylF), *O*-tert-Butyl-L-tyrosine (tBtylY), L-2-Naphthylalanine (NapA), L-4-Acetylphenylalanine (pAcF), L-4-Iodophenylalanine (pIF), L-4-Bromophenylalanine (pBrF), L-4-Chlorophenylalanine (pClF), L-4-Fluorophenylalanine (pFF), L-4-Azidophenylalanine (pAzF), L-4-Nitrophenylalanine (pNitroF), L-3-Iodophenylalanine (mIF), L-phenylalanine, L-tyrosine, L-tryptophan, and 5-Hydroxytryptophan (5OHW). Solutions of amino acids (50 or 100 mM) were made in 10-50 mM NaOH.

### Minimal Media SAA Spiking Experiments

Minimal media adapted C321.ΔA strains^49^ harboring either (i) pZE21/Ub-M-sfGFP_151UAG only, (ii) pZE21/Ub-M-sfGFP_151UAG and 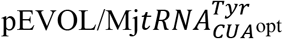, (iii) pZE21/Ub-M-sfGFP_151UAG only and pEVOL/bipARS_WT-tRNA_WT, or (iv) pZE21/Ub-M-sfGFP_151UAG only and pEVOL/bipARS_10-tRNA_10 were inoculated from frozen stocks in at least experimental duplicates. A 1X M9 salt medium containing 6.78 g/L Na_2_HPO_4_ ·7H_2_O, 3 g/L KH_2_PO_4_, 1 g/L NH_4_CI, and 0.5 g/L NaCl, supplemented with 2 mM MgSO_4_, 0.1 mM CaCl_2_, 1% glycerol, trace elements, 0.25 μg/L D-biotin, and carbenicillin was used as the culture medium. The trace element solution (100X) used contained 5 g/L EDTA, 0.83 g/L FeCl_3_ ·6H_2_O, 84 mg/L ZnCl_2_, 10 mg/L CoCl_2_ ·6H_2_O, 13 mg/L CuCl_2_O, 1.6 mg/L MnCl_2_O and 10 mg/L H_3_BO_3_ dissolved in water as previously used for metabolic engineering^50^. Inoculum were grown to confluence overnight in deep 96-well plates containing supplemented with 0.2% arabinose and chloramphenicol and/or kanamycin. Experimental cultures were inoculated at 1:7 dilution in the same media supplemented with each of the 20 standard amino acids or bipA to 1 mM or 100 uM, respectively. Cultures were incubated at 34 °C to an OD_600_ of 0.5–0.8 in a shaking plate incubator at 1050 rpm (∼4-5 h). GFP expression was induced by addition of anhydrotetracycline, and cells were incubated at 34 °C for an additional 16-20 h before measurement. All assays were performed in 96-well plate format. Cells were centrifuged at 5,000g for 5 min, washed with 1 × PBS, and resuspended in 1 × PBS after a second spin. GFP fluorescence was measured on a Biotek spectrophotometric plate reader using excitation and emission wavelengths of 485 and 525 nm. Fluorescence was then normalized by the OD_600_ reading to obtain FL/OD. Average normalized FL/OD from 3 independent experiments were plotted.

### NSAA Incorporation Assays

Strains harboring integrated GFP reporters and AARS/tRNA plasmids were inoculated from frozen stocks in biological triplicate and grown to confluence overnight in deep well plates. Experimental cultures were inoculated at 1:100 dilution in either LB^L^ or 2XYT media supplemented with chloramphenicol, arabinose, and the appropriate NSAA. Cultures were incubated at 34 °C to an OD_600_ of 0.5–0.8 in a shaking plate incubator at 400 rpm (∼4-5 h). GFP expression was induced by addition of anhydrotetracycline, and cells were incubated at 34 °C for an additional 16-20 h before measurement.

All assays were performed in 96-well plate format. Cells were centrifuged at 5,000g for 3 min, washed with PBS, and resuspended in PBS after a second spin. GFP fluorescence was measured on a Biotek spectrophotometric plate reader using excitation and emission wavelengths of 485 and 525 nm (Gain = 80). Fluorescence signals were corrected for autofluorescence as a linear function of OD_600_ using the parent C321.ΔA strain that does not contain a reporter. Fluorescence was then normalized by the OD_600_ reading to obtain FL/OD.

### Library Generation

Error-prone PCR (EP-PCR) was performed using the GeneMorph II Random Mutagenesis Kit (Stratagene Catalog #200550), following manufacturer instructions to obtain approximately an average of 2-4 DNA mutations per library member. To generate libraries of MjTyrRS-derived AARSs, roughly 175 ng of PCR template was used in each 25 uL of PCR mix containing primers that have roughly 40 base pairs of homology flanking the AARS coding region. The reaction mixture was subject to 30 cycles with Tm of 63°C and extension time of 1 min. Four separate 25 uL EP-PCR reactions were performed per AARS and then pooled. Plasmid backbone PCRs were performed using KOD Xtreme Hot Start Polymerase (Millipore Catalog #71795). Both PCR products were isolated by 1% agarose gel electrophoresis, DpnI digested, and Gibson assembled in 8 parallel 20 uL volumes per library. Assemblies were pooled, washed by ethanol precipitation, and resuspended in 50 μL of dH2O, which was drop dialyzed (EMD Millipore, Billerica, MA) and electroporated into “E. cloni” supreme cells (Lucigen, Middleton, WI). Libraries were expanded in culture and miniprepped (Qiagen, Valencia, CA). 1 μg of library was drop dialyzed and electroporated into C321.Nend for subsequent FACS experiments. Colony counts of dilutions of each transformation plated on appropriate antibiotic within one doubling time after transformation revealed library sizes of roughly 1 × 10^6^ for AARS libraries in E. cloni hosts and 1 × 10^7^ in C321.Nend hosts. 20 colonies were picked from each plate to confirm library diversity and 5-15% parent construct was observed. Transformation of Gibson assembly into E. cloni hosts was the bottleneck that determined library size, and efforts were made during all subsequent steps to ensure oversampling.

### Flow Cytometry and Cell Sorting

AARS libraries were subject to three rounds of fluorescence activated sorting in a Beckman Coulter MoFlo Astrios EQ cell sorter. Prior to each round, the NSAA incorporation assay procedure detailed above was followed such that cells would express GFP reporter proportional to the activity of the AARS library member. One notable deviation from that procedure was the use of a higher and variable inoculum volume, up to 25 uL, to avoid bottlenecking the library. Cells displaying the top 0.5% of fluorescence activation (50k cells) were collected after Round 1, expanded overnight, and used to inoculate experimental cultures for the next round. Because the next round was a negative screening round, the desired NSAA was not added into culture medium. The rest of the NSAA incorporation assay procedure was followed to eliminate cells that exhibited fluorescence due to promiscuous AARS activity on standard amino acids. In the second sort, cells displaying the lowest 10%-20% of visible fluorescence (500k cells) were collected. Cells passing the second round were expanded overnight and used to inoculate the third and final round of sorting. The experimental cultures for the third round were treated as the first round and were sorted for the upper 0.05% of fluorescence activation (1k cells). The final cells collected were expanded overnight and plated for sequencing and downstream testing. Libraries were frozen at each stage before and after sorting. FlowJo X software was used to analyze the flow cytometry data. Constructs of interest were grown overnight, miniprepped, and transformed into C321.ΔA.Ubiq-UAG-sfGFP for further analysis in plate reader assays.

### Reporter Purification

Strains harboring integrated GFP reporters and AARS/tRNA plasmids were inoculated from frozen stocks and grown to confluence overnight in 5 mL 2XYT containing chloramphenicol. Saturated cultures were used to inoculate 500 mL experimental cultures of 2XYT supplemented with chloramphenicol, arabinose, and appropriate NSAAs. Cultures were incubated at 34 °C to an OD_600_ of 0.5–0.8 in a shaking incubator at 250 rpm. GFP expression was induced by addition of anhydrotetracycline, and cells were incubated at 34 °C for an additional 24 h before measurement. Cells were centrifuged in a Sorvall RC 5C Plus at 10,000 *g* for 20 minutes. Pellets were frozen at −20 °C before lysis and purification. Lysis of resuspended pellets was performed under denaturing conditions in 10 mL 7 M urea, 0.1 M Na_2_PO_4_, 0.01 M Tris-Cl, pH 8.0 buffer with 450 units of Benzonase (Novagen, cat. no. 70664-3) using 15 minutes of sonication in ice using a QSonica Q125 sonicator. Lysate was distributed into microcentrifuge tubes and centrifuged for 20 minutes at 20,000 *g* at room temperature, and then protein-containing supernatant was removed. 2 mL supernatant with 7.5 uM imidazole was added to 250 uL Ni-NTA resin (Qiagen Cat no. 30210) and equilibrated at 4°C overnight. Columns were washed with 7x 1 mL washes using 8 M urea, 0.1 M Na_2_PO_4_, 0.01 M Tris-Cl. Wash 1 and 2 were adjusted to pH 6.3 and contained no imidazole. Washes 3-7 were adjusted to pH 6.1 and contained imidazole at concentrations of 10 mM, 25 mM, 40 mM, 60 mM and 80 mM respectively. Protein was eluted with two 150 uL elutions using elution buffer (8 M urea, 0.1 M Na_2_PO_4_, 0.01 M Tris-Cl, pH 4.5, 300 mM imidazole). Gels demonstrated that wash 5 eluted the protein, and for several samples the wash 5 fraction was concentrated ∼20X using Amicon Ultra 0.5 mL 10K spin concentrators. Protein gels were loaded with 30 uL wash or elution volumes along with 10 uL Nu-PAGE loading dye in Nu-PAGE 10% Bis-Tris Gels (ThermoFisher Cat. no NP0301). Protein gels were run at 180 V for 1 h, washed 3x with DI water, stained with coomassie (Invitrogen Cat. no LC6060) for one hour. Gels were destained overnight in water on a shaker at room temperature and images were taken with a BioRad ChemiDoc MP imaging system.

### Mass Spectrometry

Samples were submitted for single LC-MS/MS experiments that were performed on a LTQ Orbitrap Elite (Thermo Fischer) equipped with Waters (Milford, MA) NanoAcquity HPLC pump. Trypsin-digested peptides were separated onto a 100 μm inner diameter microcapillary trapping column packed first with approximately 5 cm of C18 Reprosil resin (5 μm, 100 Å, Dr. Maisch GmbH, Germany) followed by analytical column ∼20 cm of Reprosil resin (1.8 μm, 200 Å, Dr. Maisch GmbH, Germany). Separation was achieved through applying a gradient from 5–27% ACN in 0.1% formic acid over 90 min at 200 nl min–1. Electrospray ionization was enabled through applying a voltage of 2.0 kV using a home-made electrode junction at the end of the microcapillary column and sprayed from fused silica pico tips (New Objective, MA). The LTQ Orbitrap Elite was operated in the data-dependent mode for the mass spectrometry methods. The mass spectrometry survey scan was performed in the Orbitrap in the range of 395 –1,800 m/z at a resolution of 6 × 10^4^, followed by the selection of the twenty most intense ions (TOP20) for CID-MS2 fragmentation in the Ion trap using a precursor isolation width window of 2 m/z, AGC setting of 10,000, and a maximum ion accumulation of 200 ms. Singly charged ion species were not subjected to CID fragmentation. Normalized collision energy was set to 35 V and an activation time of 10 ms, AGC was set to 50,000, the maximum ion time was 200 ms. Ions in a 10 ppm m/z window around ions selected for MS2 were excluded from further selection for fragmentation for 60 s.

### Mass Spectrometry Analysis

Raw data were submitted for analysis in Proteome Discoverer 2.1.0.81 (Thermo Scientific) software. Assignment of MS/MS spectra was performed using the Sequest HT algorithm by searching the data against a user provided protein sequence database as well as all entries from the E. coli Uniprot database and other known contaminants such as human keratins and common lab contaminants. Sequest HT searches were performed using a 20 ppm precursor ion tolerance and requiring each peptides N-/C termini to adhere with Trypsin protease specificity while allowing up to two missed cleavages. Cysteine carbamidomethyl (+57.021) was set as static modifications while methionine oxidation (+15.99492 Da) was set as variable modification. MS2 spectra assignment false discovery rate (FDR) of 1% on protein level was achieved by applying the target-decoy database search. Filtering was performed using a Percolator. For quantification, a 0.02 m/z window centered on the theoretical m/z value of each the six reporter ions and the intensity of the signal closest to the theoretical m/z value was recorded. Reporter ion intensities were exported in result file of Proteome Discoverer 2.1 search engine as an excel tables. All fold changes were analyzed after normalization between samples based on total unique peptides ion signal.

### *In vitro* Aminoacylation Assays

Wild-type BipARS, BipARS9, and BipARS10 DNA template was amplified from the pEVOL.BipARS plasmid and cloned into pET20b using Gibson assembly (New England Biolabs) with primers pET20.F2 and pET20.R for linearization of pET20b and BipRS.F and BipRS.R2 for amplification of BipARS. The BipARS.pET20b plasmids were transformed into BL21(DE3) cells. A 25-mL overnight culture was used to inoculate 500 mL of fresh LB media containing ampicillin. Cells were grown at 37 °C to an OD_600_ of approximately 0.6, and protein overexpression was induced with 1 mM IPTG for 4 h. Cells were harvested by centrifugation at 4 °C for 20 minutes at 6000 rpm. Cells were lysed using 50 mM Tris (pH7.5), 300 mM NaCl, 3 mM 2-mercaptoethanol and 5 mM imidazole followed by sonication. Lysed cells were centrifuged at 18000 × g for 1 h at 4 °C. The supernatant was run through TALON resin and BipARS was eluted using an imidazole concentration gradient. The proteins were stored in 50 mM HEPES (pH 7.3), 50 mM KCl, and 1 mM dithiothreitol (DTT). Protein concentration was calculated using the Bradford assay (BioRad).

The tRNA genes were cloned into pUC18 using Gibson Assembly. pUC18 was linearized using primers pUCbip_F and pUCbip_R. The tRNA gene fragment was prepared by annealing 2 μM of primers tBip_F and tBip_R for WT tRNA, tBip9_F and tBip9_R for tRNA variant 9, and tBip10_F and tBip10_R for tRNA variant 10. tRNAs were obtained by in vitro transcription using T7 RNA polymerase. ∼100 μg of resulting plasmid was digested with BstNI overnight at 55 °C, and the digestion reaction was used to start in vitro transcription by adding transcription buffer (40 mM Tris-HCl, pH 8, 6 mM MgCl_2_, 1 mM spermidine, 0.01% Triton, 0.005 mg/mL BSA, and 5 mM dithiothreitol), 4 mM NTPs (ATP, GTP, UTP, and CTP), 20 mM MgCl_2_, 5 mM DTT, 2 units/mg of pyrophosphatase (Roche), and 0.75 mg/mL T7 RNA polymerase. The reaction was incubated for 6-7 h at 37 °C. The tRNA was purified using an 8 M urea/12 % acrylamide gel and extracted from the gel using a solution containing 0.5 M sodium acetate and 1 mM EDTA (pH 8) overnight at 30 °C followed by ethanol precipitation.

For aminoacylation reactions, tRNAs were radiolabeled at the 3’-end using CCA-adding enzyme as previously described (*51*). Reactions were carried out with 5 μM tRNA (with trace amount of ^32^P-labeled tRNA), 2.5 mM amino acid, and 5 pM BipARS in buffer containing 50 mM HEPES (pH 7.3), 4 mM ATP, 20 mM MgCh, 0.1 mg/mL BSA, and 1 mM DTT. Reactions were incubated for 30 minutes at 37 °C. 2 μL of reaction mixture were quenched in 5 μL of 0.1 U/μL P1 nuclease (Sigma) in 200 mM sodium acetate (pH 5) right after enzyme addition and after 30 min. The quenched time points were incubated at room temperature for 1 h. 1 μL of the solution was run PEI cellulose thin layer chromatography sheets. The fraction of aminoacylated tRNA was determined as described previously^51^. All assays were repeated three times. Figures were generated using Prism 7 (GraphPad Software).

### Biocontainment Escape Frequency Assays

Escape assays were performed nearly as previously described^13^. All strains were grown in permissive conditions and harvested in late exponential phase. Cells were washed twice in LB and resuspended in LB. Viable CFU were calculated from the mean and standard error of the mean (SEM) of three technical replicates of tenfold serial dilutions on permissive media. Three technical replicates were plated on non-permissive media and monitored for 7 days. Synthetic auxotrophs were plated on two different non-permissive media conditions: SCA - LB with SDS, chloramphenicol, and arabinose – for previously published strains; and KA - LB with kanamycin and arabinose – for strains generated in this study. The latter strains were isolated by transformation with pEVOL vectors harboring kanamycin resistance markers instead of chloramphenicol resistance markers. Passaging and replica plating were used to ensure that isolated strains had lost chloramphenicol resistance and thus the original OTS construct used in the previous study. If synthetic auxotrophs exhibited escape frequencies above the detection limit (lawns) on non-permissive media at days 2, 5, or 7, escape frequencies for those days were calculated from additional platings at lower density. The SEM across technical replicates of the cumulative escape frequency was calculated as previously indicated.

### Biocontained strain doubling time measurement

Doubling times for biocontained strains were measured in triplicate by plate reader as indicated earlier for growth assays. Doubling time assays for biocontained strains in the presence of only non-cognate NSAAs were performed as follows: cells grown to mid-log in permissive media were washed twice in LB and diluted to OD ∼0.1 before 300-fold dilution into three 150 μL volumes of LB+NSAA for each NSAA. These cultures were incubated in the Eon plate reader at conditions described earlier.

## Acknowledgements

We thank Dr. Daniel J. Mandell (Harvard), Dr. Ethan Garner (Harvard), Dr. Karl Schmitz (MIT), Dr. Irene M. B. Reizman (Rose-Hulman), and Bernardo Cervantes (MIT) for insightful discussions. We thank Chad Araneo for FACS assistance and Dr. Bogdan Budnik for mass spectrometry assistance. This project was graciously funded by the U.S. Dept. of Energy, grant DE-FG02-02ER63445 (GMC), and the National Institutes of Health, grants R01GM22854 and R35GM122560 (DS). GMC has related financial interests in ReadCoor, EnEvolv, and GRO Biosciences. AMK and GMC have filed a provisional patent on PTP, and AMK/DS/EK/GMC have filed a provisional patent on evolved BipA OTS variants. For a complete list of GMC’s financial interests, please visit http://arep.med.harvard.edu/gmc/tech.html. Author Contributions: AMK conceived the project and designed and performed most experiments. DAS generated EP-PCR AARS libraries, cloned AARS/tRNA combinations, purified reporter protein, and submitted samples for mass spectrometry. AMK, DAS, and EK screened AARS libraries. EK performed minimal media AA spiking experiments. OVR performed AARS and tRNA purification and biochemical characterization of tRNA aminoacylation. ML generated strains containing N-end selectable markers. DS and GMC supervised experiments. AMK and DAS led manuscript writing and all authors reviewed before submission.

## Supplementary Information

Supplementary Text

Supplementary Figures 1-11

Supplementary Tables 1-4

